# Multi-color super-resolution imaging to study human coronavirus RNA during cellular infection

**DOI:** 10.1101/2021.06.09.447760

**Authors:** Jiarui Wang, Mengting Han, Anish R. Roy, Haifeng Wang, Leonhard Möckl, Leiping Zeng, W. E. Moerner, Lei S. Qi

## Abstract

The severe acute respiratory syndrome coronavirus 2 (SARS-CoV-2) is the third human coronavirus within 20 years that gave rise to a life-threatening disease and the first to reach pandemic spread. To make therapeutic headway against current and future coronaviruses, the biology of coronavirus RNA during infection must be precisely understood. Here, we present a robust and generalizable framework combining high-throughput confocal and super-resolution microscopy imaging to study coronavirus infection at the nanoscale. Employing the model human coronavirus HCoV-229E, we specifically labeled coronavirus genomic RNA (gRNA) and double-stranded RNA (dsRNA) via multicolor RNA-immunoFISH and visualized their localization patterns within the cell. The exquisite resolution of our approach uncovers a striking spatial organization of gRNA and dsRNA into three distinct structures and enables quantitative characterization of the status of the infection after antiviral drug treatment. Our approach provides a comprehensive framework that supports investigations of coronavirus fundamental biology and therapeutic effects.

## INTRODUCTION

Severe acute respiratory syndrome coronavirus 2 (SARS-CoV-2) is the causative viral pathogen for the ongoing COVID-19 pandemic (Wu et al., 2020; Zhou et al., 2020). SARS-CoV-2 is one of the seven coronaviruses (CoVs) known to infect humans. HCoV-229E and HCoV-NL63 belong to Alphacoronaviruses; HCoV-OC43, HCoV-HKU1, SARS-CoV, MERS-CoV, and SARS-CoV2 are Betacoronaviruses (Fung and Liu, 2019; V’Kovski et al., 2021). After SARS-CoV and MERS-CoV, SARS-CoV-2 is the third human coronavirus that causes wide-spread respiratory syndromes with severe and often fatal disease progression (Cevik et al., 2021; Hu et al., 2020; Petersen et al., 2020; V’Kovski *et al*., 2021; Zhu et al., 2020).

All CoVs are enveloped, single-stranded, positive-sense RNA viruses. The genome size is around 30 kilobases (kb), making it one of the largest known RNA viral genomes (Gorbalenya et al., 2006; Kim et al., 2020). All of the viral nonstructural proteins (nsps), including components of the RNA-dependent RNA polymerase (RdRp) complex, are encoded in the two open reading frames that encompasses roughly the first two-thirds of the genome (Hartenian et al., 2020). Upon entering the cytoplasm of the host cell, this part of the genome is directly translated by the host translation machinery into polyproteins, which are cotranslationally cleaved by proteases that are part of the polyproteins (Hartenian *et al*., 2020). Located within one-third of the genome at the 3’ end are four structural proteins, common to all CoVs: the spike (S), nucleocapsid (N), membrane (M), and envelope (E) proteins. Additionally, some virus-specific accessory proteins sit between the structural proteins (Kim *et al*., 2020). The structural proteins and accessory proteins are expressed by subgenomic mRNAs (sgRNAs) (Sola et al., 2015). The RdRp complex is responsible for all viral RNA synthesis, including generating negative-sense full-length genomic RNA (gRNA) and sgRNAs from the viral gRNA template (Hilgenfeld and Peiris, 2013; Posthuma et al., 2017; Snijder et al., 2016). These negative stranded gRNA and sgRNAs are used as templates for the synthesis of the positive stranded gRNA and sgRNAs, which are the mRNAs used for the synthesis of the viral proteins (Hartenian *et al*., 2020). CoVs rely on the infrastructure of the host cell for all stages of life. Besides the translation machinery, CoVs have been shown to heavily modify and repurpose the inner membranes of the host cell to generate separate compartments, such as double membrane vesicles (DMVs) and convoluted membranes (CMs) (Knoops et al., 2008). The endoplasmic reticulum-Golgi intermediate compartment (ERGIC) is used for virion assembly before being exported from the cell through exocytosis (Klumperman et al., 1994; Stertz et al., 2007). These compartments likely allow the virus to replicate and assemble without alerting the host immune system and provide a mechanism to concentrate viral RNA, proteins, and nucleotides for optimal amplification (Hartenian *et al*., 2020).

The scientific community has gained a remarkable amount of knowledge about coronavirus biology in terms of the genomics/biochemistry, the key protein structures (from averaged single-particle cryo-electron microscopy (cryo-EM)), and the cellular invasion mechanism (from cryoelectron tomography (cryo-ET)) (Ke et al., 2020; Klein et al., 2020; Knoops *et al*., 2008; Walls et al., 2020; Wrapp et al., 2020; Zhang et al., 2020). However, various aspects of coronavirus RNA biology during infection of human cells – for example, how the different stages of the viral replication cycle are coordinated and spatially organized within the cell – are largely unknown. Cryo-EM and -ET indeed show many gray-scale structures at high resolution and a high degree of resulting context, but the molecular identity of proteins and oligonucleotides is lost (Lyumkis, 2019; Turk and Baumeister, 2020). Importantly, precise and specific fluorescence labeling lights up specific biomolecules, which can be imaged both by diffraction-limited (DL) and by super-resolution (SR) microscopy with comparably high throughput, enabling the examination of spatial localization patterns in the cell with specific cellular context and spatial resolution down to ~10 nm.

Here, we introduce a multi-color fluorescence imaging framework combining confocal and SR imaging approaches to examine the spatial interactions between viral RNA and viral factors during CoVs infection. We demonstrate the efficacy of our approach using the HCoV-229E coronavirus in MRC5 normal lung fibroblasts (Jacobs et al., 1970), concentrating on specific fluorescence labeling of two key oligonucleotide players: the viral gRNA, and double stranded RNA (dsRNA). We present a model for HCoV-229E RNA spatial organization during infection that incorporates our observations of three different specific structures: large gRNA clusters, very tiny nanoscale gRNA puncta containing a single copy of the genome, and round intermediate-sized puncta highlighted by the dsRNA label. While multi-color confocal imaging allows us to rapidly screen across many cells during the time course of infection, our SR images illustrate the spatial relations of viral gRNA and the endoplasmic reticulum (ER), dsRNA and ER, gRNA and the spike protein, gRNA and total positive-sense viral RNAs ((+) vRNA), as well as gRNA and dsRNA at the nanoscale providing complementary information to previous electron microscopy studies. In particular, our work demonstrates a striking spatial separation between dsRNA and gRNA in contrast to previous observations, highlighting unusual aspects of the spatial organization within the modified inner membrane of ER. Moreover, Remdesivir, a nucleoside analogue that perturbs viral RNA replication (Agostini et al., 2018; Beigel et al., 2020; Spinner et al., 2020; Wang et al., 2020), decreased the abundance of both dsRNA and gRNA without decreasing the sizes of dsRNA puncta, providing additional evidence for the active site of RNA replication. Our framework provides an imaging-based system to study the pathogenesis of current and emerging coronaviruses, and their interaction with antiviral drugs. It will aid in quantitatively dissecting different stages of the human coronavirus life cycle and in revealing functional virus-host interactions. Due to the robustness and ease of implementation of our framework, the approach will benefit both fundamental investigations of human coronavirus biology as well as potential therapeutic interventions, helping to address a key challenge in human health.

## RESULTS

### Confocal imaging of coronavirus gRNA and dsRNA

It has been proposed that coronaviruses replicate their gRNA in association with the ER (Harcourt et al., 2004; Ivanov et al., 2004; Knoops *et al*., 2008; Prentice et al., 2004; Snijder et al., 2006). To study the spatial localization of viral gRNA of HCoV-229E with respect to the ER membrane, we first performed two-color confocal imaging of gRNA and ER in MRC5 cells. We designed and synthesized a set of 48 fluorescence *in situ* hybridization (FISH) probes, specifically targeting the RdRp-coding region (from here on referred to as the gRNA FISH probes), which exclusively recognize gRNA and not sgRNAs (**Figure 1A**) (Marra et al., 2003; Viehweger et al., 2019). To study the localization of dsRNAs, we used a well-known anti-dsRNA antibody that recognizes dsRNAs longer than 40 base pairs (Schonborn et al., 1991) and a corresponding CF568-labeled secondary antibody for immunostaining (**Figure 1B**). The ER was fluorescently labeled in MRC5 cells using lentivirus transduction through genetic encoding of Sec61B, a single-pass component of the channel-forming translocon complex on ER (Lang et al., 2017), fused with a green fluorescent protein (GFP) from *Aequorea coerulescens.* (**Figure 1C**). (All subsequent experiments not involving ER were performed on wild-type cells without transduction.)

**Figure 1.**
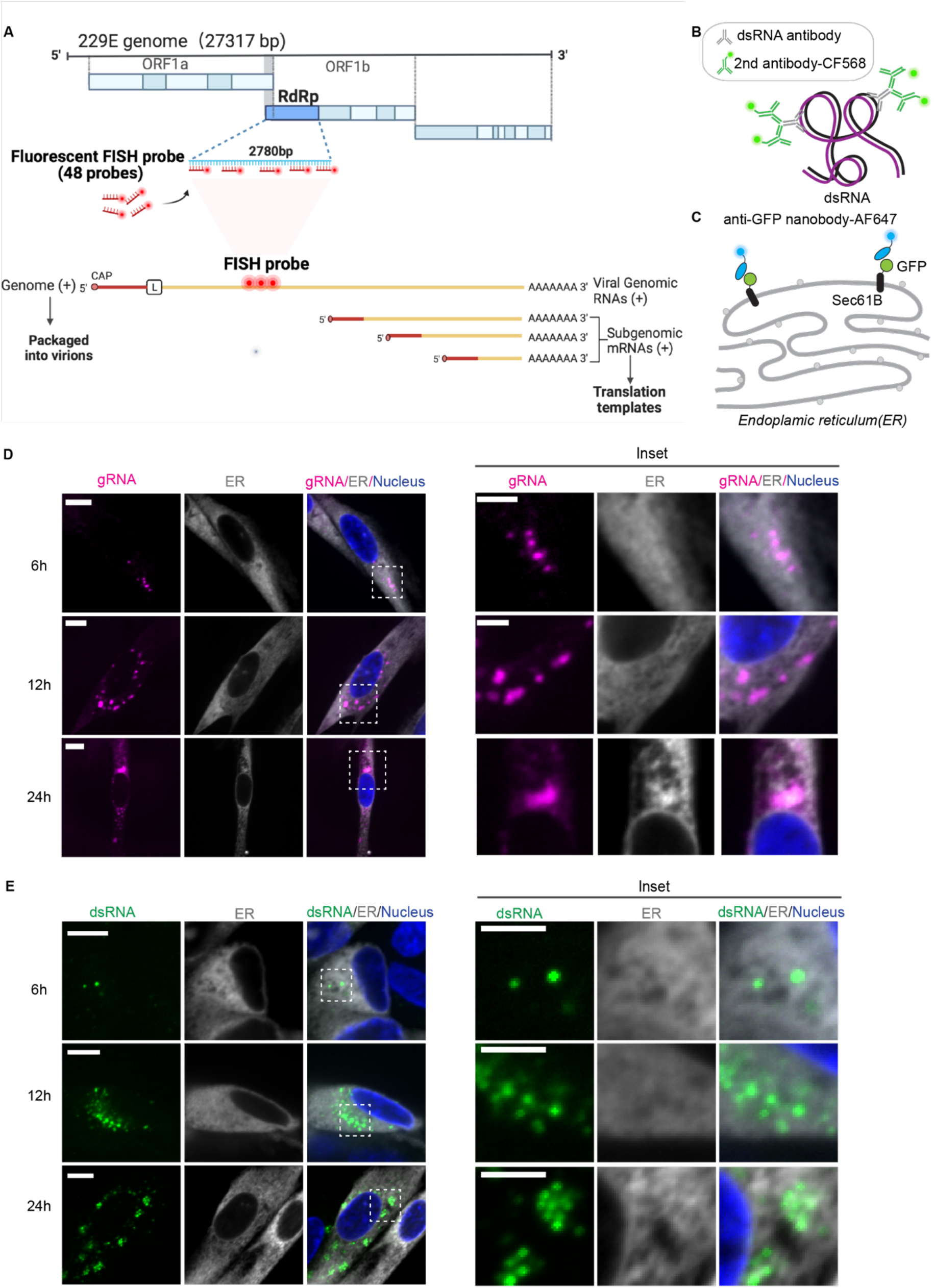
Visualization of the positive-sense viral gRNA and dsRNA. **(A)** Top: 229E genomic construct map used for the detection of viral genome in fixed cells. 48 FISH probes were designed to target genomic RNA (gRNA) by hybridizing with the RNA-dependent RNA polymerase (RdRp)-coding region, which is only present in the positive-sense gRNA and not in the subgenomic RNAs (sgRNAs). Bottom: Viral genome compared to sgRNAs, which do not contain complementary sequences to the FISH probes. The FISH probes are labeled by CF568 or AF647. Adapted from templates “Discontinuous Transcription” and “Remdesivir Active Molecule Interaction with SARS-CoV-2 RdRp”, by BioRender.com (2021). Retrieved from https://app.biorender.com/biorender-templates. (Hartenian *et al*., 2020) **(B)** Scheme showing double-stranded RNA (dsRNA) targeted by anti-dsRNA antibodies in fixed cells. Each anti-dsRNA antibody recognizes 40 bp of dsRNA. A CF568-labeled secondary antibody is then introduced for visualization. **(C)** Scheme showing the endoplasmic reticulum (ER) visualized in fixed cells via the expression of a single-pass ER membrane protein, GFP-Sec61B. An AF647-labeled anti-GFP nanobody is used to perform super-resolution (SR) imaging of the ER. **(D)** Representative confocal images of gRNA (magenta) in GFP-Sec61B-expressing fixed MRC5 cells (gray). Cells were infected with 0.2 MOI 229E and fixed at 6-, 12-, and 24-hour post infection (h p.i.). Nucleus is labeled with DAPI (blue). Scale bar: 10 μm. Right: insets of dotted boxes. Scale bar: 5 μm. **(E)** Representative confocal images of dsRNA (green) in GFP-Sec61 B-expressing fixed MRC5 cells (gray) at 6, 12, and 24 h p.i. with 0.2 MOI 229E. The nucleus is labeled with DAPI (blue). Scale bar: 10 μm. Right: insets of dotted boxes. Scale bar: 5 μm.

We examined the localization of gRNA at different time points after HCoV-229E infection by conventional confocal microscopy (**Figure 1D**). We applied 0.2 multiplicity of infection (MOI) of virus to MRC5 cells, which were fixed at various time points before labeling. At 6 h p.i., we detected distinct clusters of gRNA in the cytoplasmic region of the cells. At 12 or 24 h p.i., we observed more and larger clusters of gRNA concentrated at the perinuclear regions. Some large gRNA clusters showed colocalization with Sec61B-GFP (e.g., **Figure 1D**, 24 h p.i.) at the DL resolution of confocal microscopy.

Several studies suggest that the replication and transcription of coronaviruses take place within DMVs and/or convoluted membranes CMs (Kirkegaard and Jackson, 2005; Kopek et al., 2007; Miller and Krijnse-Locker, 2008; Netherton et al., 2007). For example, EM studies showed that DMVs contain dsRNAs, which are intermediates of coronavirus replication and transcription(Knoops *et al*., 2008; Schonborn *et al*., 1991). We observed small puncta of dsRNA clusters at 6 h p.i., and more dot-shaped puncta at 12 and 24 h p.i. These puncta might represent the cellular localization of active transcription of HCoV-229E coronavirus or merely that of the transcription/replication intermediates **(Figure 1E).**

### SR microscopy shows distinct spatial arrangements of dsRNA and gRNA relative to the ER

While confocal microscopy provides a high-throughput approach to screen many samples during infection, its optical DL resolution (approximately 300 nm) prevents us from accessing the length scales relevant to the identification of the precise spatial arrangement between the ER and gRNA or dsRNA, and clearly observing completed virions which only have a single copy of the gRNA.

To map the spatial relationship between the ER membrane and HCoV-229E beyond the diffraction limit, we performed two-color SR imaging by staining the ER and dsRNA with AF647 labeled anti-GFP nanobody and CF568 conjugated secondary antibody against dsRNA primary antibody. To avoid bleaching of AF647 via 561 nm irradiation, AF647 was imaged first, followed by CF568. All cells were fixed prior to imaging.

The SR reconstructions clearly resolved individual circularly shaped dsRNA puncta in the context of the ER network, allowing a precise determination of their spatial relation (localization precision σ×,y ~10 nm, **Supplementary Figure 2F**). Numerous puncta appeared to be adjacent to the ER membrane, suggesting possible membrane connection between the two. In addition, many dsRNA puncta (green) were completely separated from the ER membrane (gray), appearing in voids of the ER network (**Figure 2A-B and Supplementary Figure 2A-B**). Some dsRNA clusters were as far as approximately 500 nm away from the closest continuous ER membrane, a substantial distance considering the high density of the ER. Therefore, we postulate that these distant, isolated and round dsRNA puncta are surrounded by membranes, as in a DMV, which are severed from the ER network. Interestingly, the lack of a ring-like ER signal surrounding dsRNA puncta suggests that membranes modified by viruses do not contain Sec61B and potentially other components normally present in the ER, but this may be due to the relatively high curvature of a DMV surface preventing single-pass attachment (see **Discussion**) (Derganc and Copic, 2016; Larsen et al., 2020).

**Figure 2.**
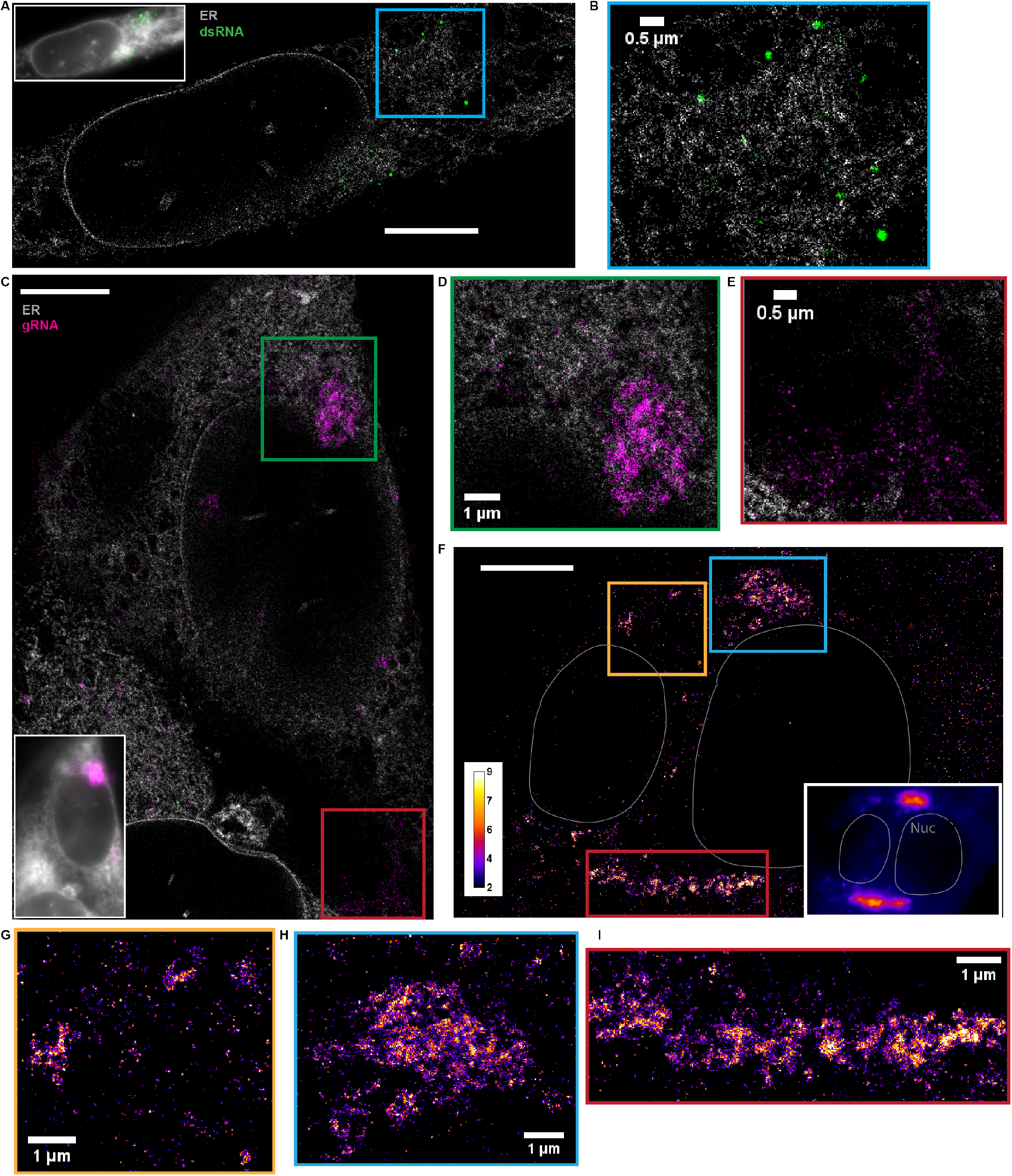
dsRNA forms puncta separated from ER while gRNA clusters frequently associate with the ER. **(A)** Two-color SR reconstruction of a cell at 24 h p.i. where the ER (gray) and dsRNAs (green) are labeled. Scale bar: 5 μm. **(B)** Zoom-in of the boxed region. dsRNA forms compact puncta that appear in regions outside of the ER network. **(C)** Two-color SR reconstruction of a cell where the ER (gray) and gRNA (magenta) are labeled. Scale bar: 5 μm. **(D)** Zoom-ins of the green boxed region. gRNA forms clusters of various sizes, some of which exhibit poorly defined boundaries. The extended network of clusters is often closely associated with the ER membrane. **(E)** A population of the genomic RNA forms small puncta of similar sizes which are not associated with the ER (cf. Figure 3). **(F)** One-color SR reconstruction of gRNA at 12 h p.i. Inset: DL image of the same field of view. Scale bar: 5 μm. Color bar indicates the number of single-molecule detections within each pixel. **(G-I)** Zoom-ins of the colored boxed regions. Large clusters of the viral genome appear perinuclear and extend up to a few microns.

Next, we performed two-color SR imaging by staining the ER and gRNA with AF647 labeled anti-GFP nanobody and CF568-labeled FISH probes, respectively. The SR reconstructions revealed intricate gRNA structures (**Figure 2C-E and Supplementary Figure 2C-E**). We observed two key phenotypes: (i) separated nanoscale gRNA puncta that are smaller than 100 nm but larger than the achieved localization precision of 10 nm, in areas completely devoid of ER signal, and (ii) extended networks of gRNA clusters with a wide range of sizes and shapes (up to microns), often associated with the ER membrane. The shape of these extended clusters has similarities with the ER and is reminiscent of the convoluted ER membrane found in HCoV-229E infected cells (Knoops *et al*., 2008; Snijder et al., 2020).

As we suspected that the observed phenotypes change in the course of the infection due to increased viral burden, we examined gRNA distributions at 6, 12, and 24 h p.i. (**Figure 2F-I and Supplementary Figure 4A-G**) using SR imaging. We found that nanoscale gRNA puncta were present in all three time points throughout the cells. In addition, networks of larger clusters appear to grow as the infection time increases, which we further analyze below.

### SR microscopy resolves nanoscale gRNA puncta, consistent with individual gRNA particles, while large gRNA clusters can be quantified

The super-resolution reconstructions also revealed nanoscale gRNA puncta that were not observed in confocal images of the cells (**Figure 3A**). To determine the nature of the nanoscale gRNA puncta, we purified mature, packaged virions secreted from HCoV-229E infected MRC5 cells. The virions were immobilized on coverslips and labeled with the same FISH protocol that was employed for cellular labeling. SR reconstructions of the purified virions exhibited striking similarity to the nanoscale gRNA puncta observed in cells, implying that these cellular nanoscale gRNA puncta contain only one copy of the gRNA, similar to that in a packed virion. (**Figure 3A-B**). The control condition without virions verified the exquisite specificity of the FISH probes, evident from minimal nonspecific background (**Figure 3D**).

**Figure 3.**
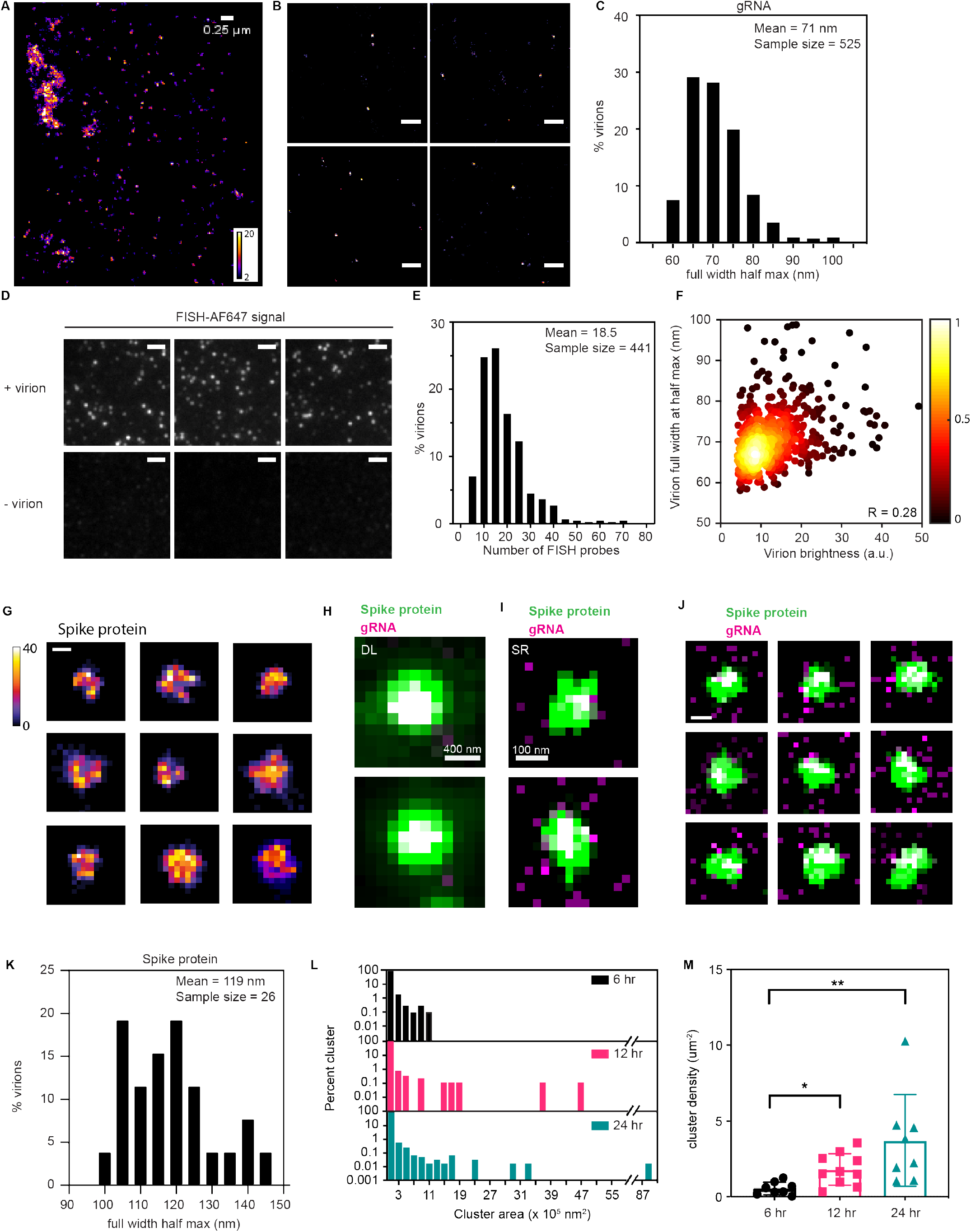
Analysis of individual virions via FISH-based labeling and SR microscopy. **(A)** Spatial distribution of fluorescently labeled gRNA in an infected cell via SR microscopy. Numerous nanoscale gRNA puncta, presumably packaged virions, appeared sometimes near the extended clusters. **(B)** SR reconstruction of isolated virions plated on a coverslip, extracted from cells labeled with the same FISH protocol employed for cellular imaging. Their appearance is very similar to the observed nanoscale gRNA puncta in the cell. Scale bar: 500 nm. **(C)** Size distribution of purified virions, calculated by fitting the virions with a 2D Gaussian. The mean full width half max of the purified virions was 71 nm. N = 525. **(D)** DL images of purified virions, immobilized on glass coverslips. FISH probes specifically targeted the viral genome with minimal nonspecific background, as evident from the control condition without virions. Scale bar: 2 μm. **(E)** Number of FISH probes per virion, calculated by dividing the total fluorescence intensity of individual virions by the singlemolecule brightness. On average, each virion is labeled by 18.5 FISH probes. N=441. **(F)** Plot of virion sizes versus brightness, colored according to the normalized local density of points. There is no clear correlation between the two parameters (Pearson correlation = 0.28). **(G)** SR reconstructions of nine unique virions stained with spike protein antibodies. The virions are 100-150 nm diameter large bright objects. Scale bar: 100 nm. **(H)** DL images of two different virions labeled with spike protein (green) and gRNA (magenta). The white color indicates clear colocalization in the center. **(I)** Two-color SR reconstructions of virions shown in panel H. A distinctive concentric structure of gRNA encapsulated by a larger structure labeled with the spike protein is clearly observed **(J)** Additional two-color SR reconstructions of several virions where colocalization is evident. 39% of the total virions observed (N=168) exhibited some degree of colocalization. Scale bar: 100 nm. **(K)** Size distributions of purified virions stained with spike protein antibody. The mean full width half maximum is 119 nm. N=26. **(L)** Percentage area distribution of gRNA clusters for cells 6 (black), 12 (magenta) and 24 (turquoise) h p.i. Cells infected for a longer period of time showed larger clusters representing increasing copy number of the viral genome. Data collected from 9, 10 and 8 cells, respectively. **(M)** Cluster density for cells 6 (black), 12 (magenta) and 24 (turquoise) h p.i. Cells infected for a longer period of time exhibited a higher cluster density. *, p< 10^-1^; **, p < 10^-2^ (two-tailed t-test). Data collected from 9, 10 and 8 cells, respectively.

To quantify the size of the virions, we fitted the SR reconstructions of the purified virions with two-dimensional Gaussians, after subjecting the reconstructions to a 1-pixel gaussian blur to reduce the contribution of pixilation due to binning. The average full width half max of 525 virions was 71 nm (**Figure 3C**), roughly consistent with previous cryo-EM studies (Barcena et al., 2009; Neuman et al., 2006). In addition, each virion was labeled with 18.5 FISH probes on average (**Figure 3E and Methods**). As the dye conjugation reaction for FISH probes had an average 75% labeling efficiency, we can conclude that most virions carried a single copy of genomic RNA. The brightness of the labeled virions did not correlate with the size of the virions, suggesting the absence of merged virions in this experiment (**Figure 3F**). Both the size and brightness of these purified virions match well with the nanoscale gRNA puncta found in cells. In combination with the fact that they were well separated from each other, our SR images suggest that the nature of these nanoscale cellular gRNA puncta contain one copy of the gRNA. Occasionally, we see higher density of such nanoscale gRNA puncta in a group (bottom left of **Figure 3A**), which might represent vesicle packets shown in previous EM studies, where multiple virions are harbored (Knoops *et al*., 2008; Ogando et al., 2020). To better understand the nature of these gRNA puncta, we performed two-color imaging experiments of gRNA and the spike protein, two biomolecules that should colocalize in a virion. This experiment also serves as a positive control for our two-color imaging method. The spike protein was labeled with a primary and secondary antibody attached to AF647 while gRNA was labeled with FISH probes labeled with CF568. Our labeling approach has very low non-specific binding (**Supplemental Figure 3A and Supplemental Figure 3C**), and DL imaging confirms that gRNA and spike colocalize in the purified virions (**Supplemental Figure 3B and Figure 3H**). For purified virions imaged with SR *in vitro,* spike labeled virions appear as approximately 100-150 nm bright round objects (**Figure 3G**). The variable shapes suggest partially formed or disassembled viral shells. Two-color SR imaging reveals that the spike protein nicely forms an outer shell with gRNA in the center, inside the shell (**Figure 3I and Figure 3J)**. Given that the spike protein is membrane embedded and labeled with ~10 nm diameter antibodies on the periphery, we expected that gRNA puncta would have a smaller footprint than the spike proteins. Quantification of the size of object using the spike label localizations (**Methods**) yields an average full width half maximum of 120 nm (**Figure 3K**), nearly 50 nm larger than that found for the SR image size based on imaging gRNA (**Figure 3C**). This concentric structure can be quantified only by the SR reconstructions. Importantly, our two-color SR images of gRNA and spike proteins of infected cells showed that the nanoscale gRNA puncta are mostly not surrounded by spike proteins, except in a very few rare cases (**Supplementary Figure 3D, orange inset**). Sometimes, gRNA localizes with spike but without being surrounded by spike (**Supplementary Figure 3D, red inset**). We conclude that the vast majority of the 70 nm gRNA puncta in the cytoplasm are likely not packaged virions prior to exocytosis, but rather gRNA alone.

We next quantified the sizes of larger gRNA structures that are more irregular in shape, possibly remodeled ER membrane (see **Discussion**). Single molecules from the SR imaging were clustered by Voronoi tessellation (**Methods and Supplement**) and the size of each cluster was determined by the summation of all areas of Voronoi polygons within the cluster (**Supplementary Figure 4H-K**) (Khater et al., 2020; Levet et al., 2015; Nicovich et al., 2017). We investigated the distribution of the sizes of gRNA clusters at 6, 12, and 24 h p.i. and observed that larger clusters formed as the infection progressed (**Figure 3L**). At 24 h p.i., some clusters grew up to approx. 9 square microns. These large clusters appear at many locations but appear more concentrated in perinuclear regions of the cells. Although we investigated three time points post infection, we observed both advanced and earlier stages of infection in each sample, leading to considerable heterogeneity among the cells. Nevertheless, the density of clusters increased as the cells were infected for a longer period (**Figure 3M**). In addition, the viral density within each cluster was relatively constant as the cluster molecule number directly correlated with cluster area with an R of 0.99 (**Supplementary Figure 4K**).

### Simultaneous visualization of gRNA and dsRNA in HCoV-229E infected cells

As SR images showed distinct spatial localization patterns between gRNA and dsRNA relative to the ER (**Figure 2**), we next investigated whether the gRNA clusters colocalize with the dsRNA puncta. We co-stained HCoV-229E-infected MRC5 cells with the FISH probes targeting gRNA and the anti-dsRNA antibody and examined many samples via confocal imaging (**Figure 4**). At 6 h p.i., only ~9% of MRC5 cells showed evident HCoV-229E gRNA and dsRNA signal (**Figure 4A&4B**). Both HCoV-229E gRNA and dsRNA appeared as dot-shaped structures, which often colocalized with each other, yet at times did not colocalize as well (see magenta and green regions which are not white), with considerable heterogeneity. At 12 and 24 h p.i., dsRNA signals were detected in the same region as genomic RNAs and formed small dot-like puncta on top of or surrounding the large clusters of gRNAs (**Figure 4A&4B**).

**Figure 4.**
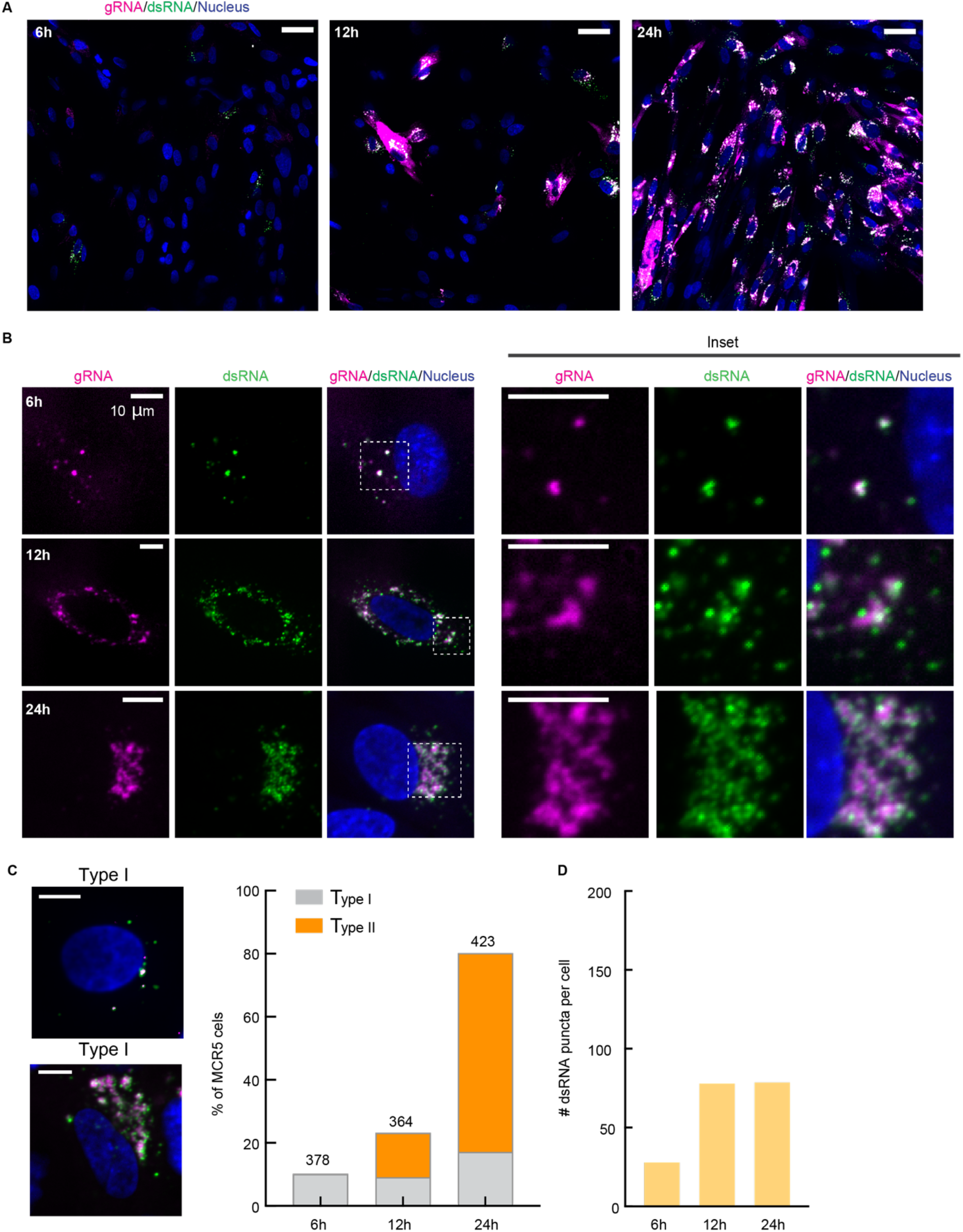
Quantification of the infection process via two-color confocal imaging of gRNA and dsRNA. **(A-B)** Representative confocal images showing gRNA (magenta) and dsRNA (green) in MRC5 cells at 6, 12, and 24 h p.i. Blue: nucleus staining. **A** shows merged images of a large view containing multiple cells using the same brightness and contrast threshold. Scale bars: 50 μm. **B** shows individual channels of representative cells and insets of selected regions (dotted squares in merged channels). Scale bars: 10 μm. **(C)** Example images (left) and quantification (right) of two types of HCoV-229E infected cells at the investigated timepoints. The number of total counted cells is listed on top of the bars. Scale bars: 10 μm. **(D)** Quantification of the number of dsRNA puncta in individual MRC5 cells at different times after HCoV-229E infection. ****, p< 10^-4^ (two-tailed t-test). Data collected from 33, 29, and 29 cells, respectively.

According to the morphology of gRNA/dsRNA clusters, we approximately classified the HCoV-229E-infected cells into two types. Type I contains localization of mostly small, scattered dot-shaped gRNA and dsRNA, which are more present at early infection. In contrast, type II shows large clusters of gRNA decorated with dot-shaped dsRNA puncta, which are more present at later infection (**Figure 4C**). We quantified the amount of dsRNA puncta per cell at different time points post infection using the same parameter threshold across all images (**Figure 4D**). While the amount of dsRNA puncta per cell present at 12 and 24 h p.i. were similar, both were significantly higher than at 6 h p.i.

As the size of many dsRNA puncta and gRNA clusters is below the diffraction limit, we turned to SR microscopy to determine the spatial correlation of the two species of viral RNAs.

### SR microscopy reveals spatial anticorrelation between gRNA clusters and dsRNA

To investigate the spatial relationship between gRNA clusters and dsRNA, we performed two-color SR imaging. In our SR reconstructions of wild type cells, as in the ER imaging of transduced cells above, dsRNA puncta are compact and circular and much smaller in size compared to the large extended gRNA clusters. Strikingly, our data clearly revealed that the dsRNA and gRNA were spatially anticorrelated across all investigated time points (**Figure 5A-D and Supplementary Figure 5-6**). dsRNA puncta generally appeared at the periphery of the extended and complex structures of the gRNA, and more distal and isolated.

**Figure 5.**
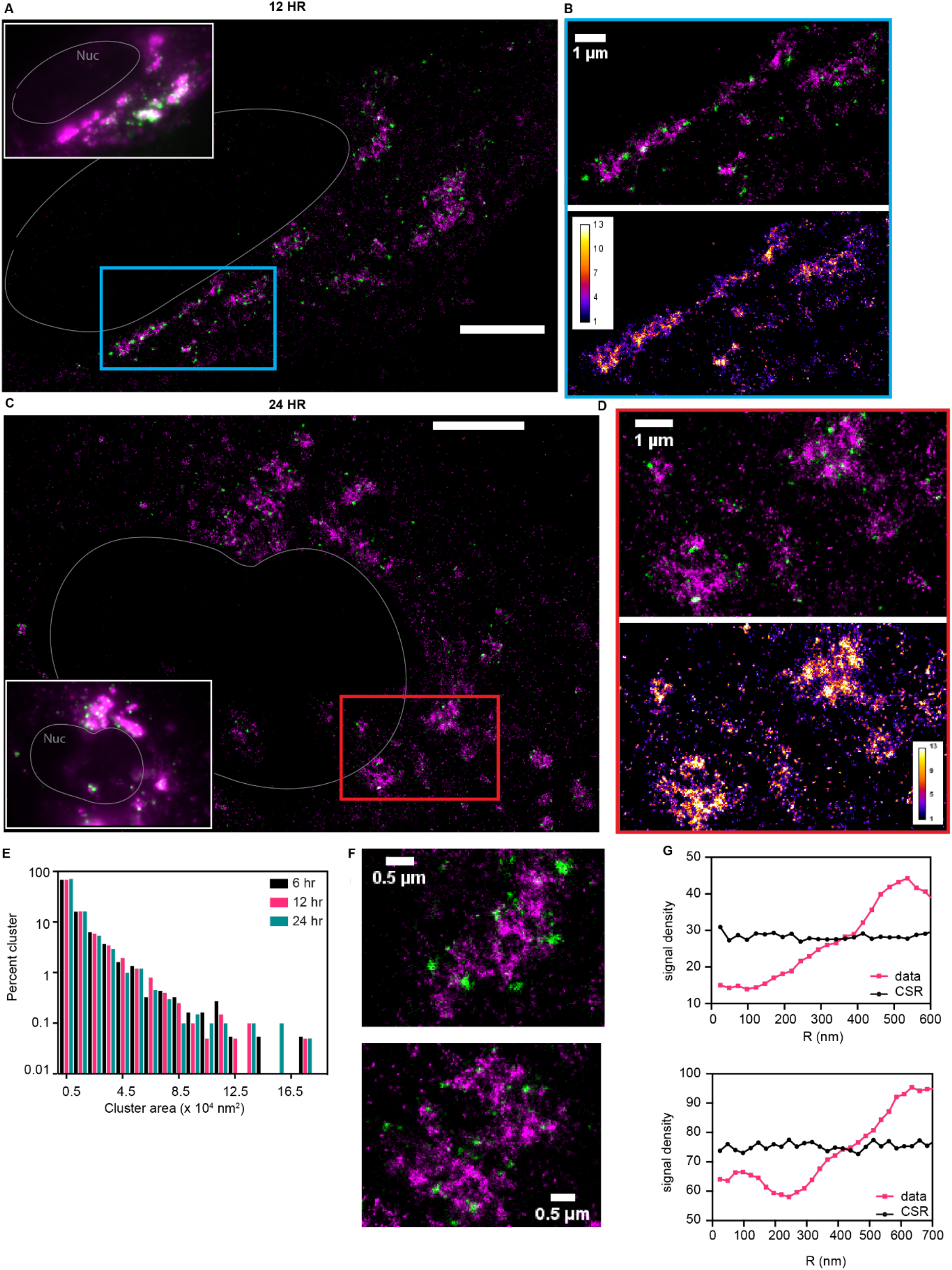
dsRNA and gRNA show anticorrelation at high resolution. **(A)** Two-color SR reconstruction of a cell 12 h p.i. gRNA (magenta) and dsRNA (green) are labeled. dsRNA signals decorate the periphery of extended gRNA clusters. Scale bars: 5 μm. **(B)** Zoom-ins of the boxed regions. The color map indicates the number of detection events per pixel for the gRNA signal. dsRNA formed compact puncta that are in the vicinity of the gRNA clusters, but spatially separated. **(C)** Two-color SR reconstruction of a cell 24 h p.i. gRNA (magenta) and dsRNA (green) are labeled. Scale bars: 5 μm. **(D)** Zoom-ins of the boxed regions. Color map as in (B). **(E)** Percentage area distribution of the dsRNA puncta at 6 (black), 12 (magenta) and 24 (turquoise) h p.i. Cells from three time points produced a similar distribution of puncta sizes. Data collected from 4 cells for each time point. The dsRNA puncta did not grow beyond 400 nm in diameter. **(F-G)** Spatial anticorrelation between the gRNA clusters (magenta) and dsRNA puncta (green) for the shown fields of view, quantified via spatial point statistics. Complete spatial randomness as a reference was simulated with the same signal density (black). Scale bar: 250 nm.

To quantitatively characterize the spatial correlation between dsRNA puncta close to gRNA clusters, we performed analysis based on spatial point statistics as previously described (Bayas et al., 2018). Briefly, for each localization event in one channel, the number of localization events in the other channel is extracted for various distances. For complete spatial randomness (CSR), this approach yields a constant signal density for any radius. Deviation from CSR allows the identification of characteristic spatial relations between the two channels. In our case, we could clearly verify the strong spatial anticorrelation already visible by eye. In addition, our quantitative analysis indicates that gRNA clusters are surrounded by dsRNA puncta at 50 – 300 nm distances (**Figure 5F-G**). Individual dsRNA puncta could be as far as up to 400 – 500 nm away from the closest gRNA cluster. This result underscores the fact that dsRNA detection is orthogonal to our FISH probe labeling. This could either be due to the inaccessibility of binding sites for the FISH probe when the strands are hybridized, competing binding between FISH probes and the dsRNA antibodies, or the unlikely scenario that the dsRNA does not contain positive sense genomic RNA but rather sgRNAs.

Our two-color imaging workflow detects anti-correlation (dsRNA and gRNA) as well as correlation (spike and gRNA in a purified virion, **Figure 3**). Given that correlation was observed *in vitro,* we wanted to confirm that correlation can be observed in cells. Accordingly, another set of FISH probes, (+) vRNA FISH probes, were designed to target the N-protein coding region and 3’-UTR, a label which binds to all (+) vRNAs including full length gRNA as well as sgRNAs (**Supplementary Figure 1 and Supplementary Figure 7**). FISH probes targeting gRNA and total RNAs are orthogonal as demonstrated by our controls (**Supplementary Figure 7A**). When we performed two-color SR imaging of gRNA and (+) vRNAs, we observed regions with colocalization as well as areas without colocalization. Extended gRNA clusters appear to be labeled with both FISH probes, indicated by colocalization and white pixels (**Supplementary Figure 7B**). These results confirm our ability to observe correlation in an *in situ* system. On the other hand, sgRNAs without the RdRp coding region would only hybridize with (+) vRNA FISH probes targeting the N-protein coding region and 3’-UTR. As expected, we observed green signal from RNAs with N-protein gene and 3’-UTR only spreading across the entire cytosol. In some cases, we observed gRNA signal alone without colocalization with (+) vRNA signal. There are several possible causes for this observation. First, RNA replication proceeds from 5’ to 3’, and RNA that are being made might not contain the N-protein coding region located at the 3’ end or the 3’UTR. Second, the RdRp and the N-protein coding regions along with the 3’UTR are about 10k base pairs apart. Depending on the packaging of the RNA molecule, they could be far away from each other.

In addition, we analyzed the dsRNA puncta using Voronoi tessellation (**Methods and Supplementary Figure 4H-K**) and identified the size and molecule numbers in each punctum. Interestingly, the size distribution of dsRNA puncta remained the same across samples at 6, 12 and 24 h p.i. (**Figure 5E**). Several cryo-EM studies of various viruses showed that the diameter of DMVs could range from 100 – 400 nm (Klein *et al*., 2020; Knoops *et al*., 2008; Wolff et al., 2020). Our data indicates that the maximum radius of dsRNA puncta is up to 230 nm for HCoV-229E, consistent with the length scale of a lipid vesicle boundary, suggesting that the dsRNA is encapsulated. In addition, since we observed a wide range of dsRNA puncta sizes, different DMVs appear to be filled with dsRNA to varying degrees.

### The multi-color super-resolution imaging framework reveals coronavirus RNA changes under drug treatment

We probed the effect of Remdesivir, a nucleoside analog that inhibits RdRp activity and thus viral replication, on HCoV-229E gRNA clusters and dsRNA puncta in MRC5 cells (Agostini *et al*., 2018; Beigel *et al*., 2020; Spinner *et al*., 2020; Wang *et al*., 2020). As expected, we observed reduced percentage of MRC5 cells infected with HCoV-229E, from 100% to 30% with 0.5 μM Remdesivir (**Figure 6A**). Most cells (98%) remained infected with 0.1 μM Remdesivir treatment. However, these cells had much lower signal in both gRNA and dsRNA channels when compared with cells treated with DMSO (**Figure 6A**). In individual cells, while the relative localization between gRNA and dsRNA remained unchanged, the number of dsRNA puncta per cell significantly decreased after Remdesivir treatment (**Figure 6B**), suggesting that at least some fraction of the dsRNA is an intermediate in the viral replication process and is involved in gRNA production.

**Figure 6.**
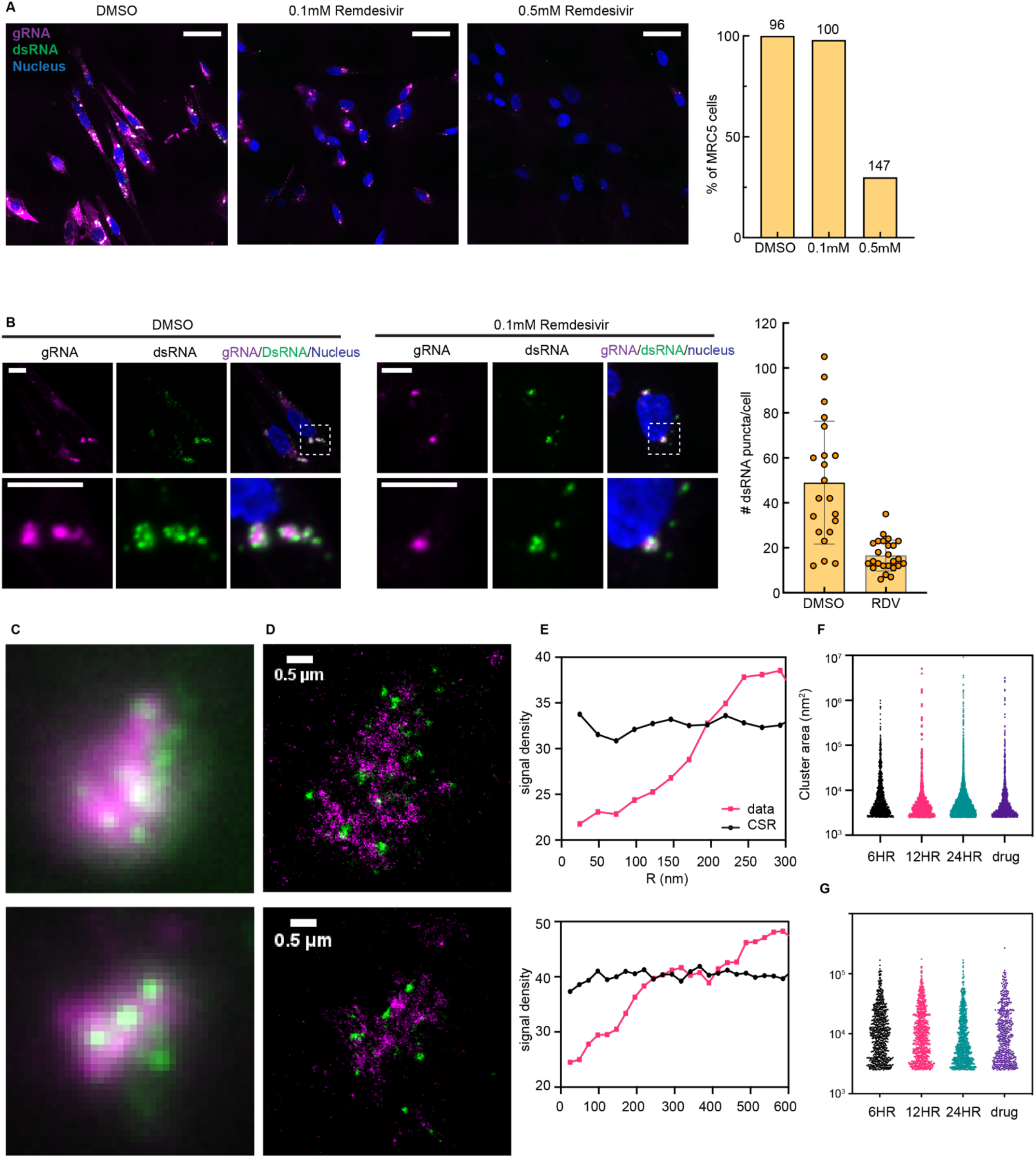
Remdesivir reduces gRNA and dsRNA content. **(A)** Representative confocal images of HCoV-229E-infected MRC5 cells in control and Remdesivir (0.1 μM and 0.5 μM) treated cells using the same brightness and contrast threshold, labeled with FISH probes targeting gRNA (magenta), anti-dsRNA antibody (green) and nuclear staining (blue). Scale bar: 50 μm. **(B)** Representative confocal images showing gRNA (magenta) and dsRNA (green) in control and 0.1μM Remdesivir-treated MRC5 cells at 24 h p.i. Right: Quantification of the number of dsRNA puncta per cell in control and 0.1μM Remdesivir-treated (RDV) MRC5 cells at 24 h p.i. Scale bar: 10 μm. ****, p< 10^-4^. Data collected from 21 and 25 cells, respectively. **(C-D)** DL images and corresponding SR reconstructions of two regions of a cell incubated with 0.1 μM Remdesivir at 24 h p.i. where gRNA (magenta) and dsRNA (green) are labeled. dsRNA puncta appeared at the periphery of gRNA clusters, again anticorrelated. **(E)** Spatial point statistics verify anticorrelation of gRNA and dsRNA. CSR is simulated with the same signal density (black). **(F)** Sizes of gRNA clusters for Remdesivir-treated cells and untreated cells. Remdesivir treatment reduced the size of the gRNA clusters. *, p< 10^-1^ (two-tailed t-test). **(G)** Sizes of dsRNA puncta for drug-treated cells and untreated cells. **, p< 10^-2^ (two-tailed t-test).

We next performed two-color SR imaging of cells stained for dsRNA and gRNA at 24 h p.i. in the presence of 0.1 μM Remdesivir. Colocalizing signals observed under DL imaging (**Figure 6C**) again turned out to be separated in the super-resolution reconstructions (**Figure 6D**). The anticorrelation of gRNA and dsRNA did not change due to Remdesivir treatment, and dsRNA puncta remained at the periphery of gRNA clusters with a wide range of distances (**Figure 6E**). As Remdesivir interferes with RdRp activity, we observed smaller gRNA clusters in cells treated with Remdesivir. This result shows that less gRNA was produced in these cells. Interestingly, the size distribution of dsRNA puncta moderately increased, to be discussed below.

## DISCUSSION

Cryo-EM and cryo-ET have revealed the intricate inner structures of coronaviruses including SARS-CoV and SARS-CoV-2 in host cells (Ke *et al*., 2020; Klein *et al*., 2020; Knoops *et al*., 2008; Walls *et al*., 2020; Wrapp *et al*., 2020; Zhang *et al*., 2020). While these very high-resolution imaging methods have illuminated the life cycle of coronaviruses, the molecular identity of the viral or host proteins and genomes is largely lost, leaving various key questions unanswered, mostly due to the lack of information on the molecular identity and/or insufficient sampling of the imaged species. Here, we developed a multi-color and multi-scale fluorescence imaging framework to visualize spatial interactions between viral RNAs and host cellular compartments at different stages of viral infection. We chose one of the seven human coronaviruses that is globally extant, HCoV-229E, in MRC5 lung cells as our model system. Despite the fact that the Alphacoronavirus HCoV-229E enters the cell via a different receptor (APN) (Yeager et al., 1992) from the Betacoronavirus, SARS-CoV2 (ACE2) (Wan et al., 2020), and from the Alphacoronavirus hCoV-NL63 (Hofmann et al., 2005), their mechanisms of replication and interactions with host machinery may considerably overlap.

We present a model for HCoV-229E RNA spatial organization during infection that incorporates our observations (**Figure 7**) of three different specific structures: the large gRNA clusters, the very tiny nanoscale puncta labeled by our gRNA label, and the round intermediate-sized puncta highlighted by the dsRNA label.

**Figure 7.**
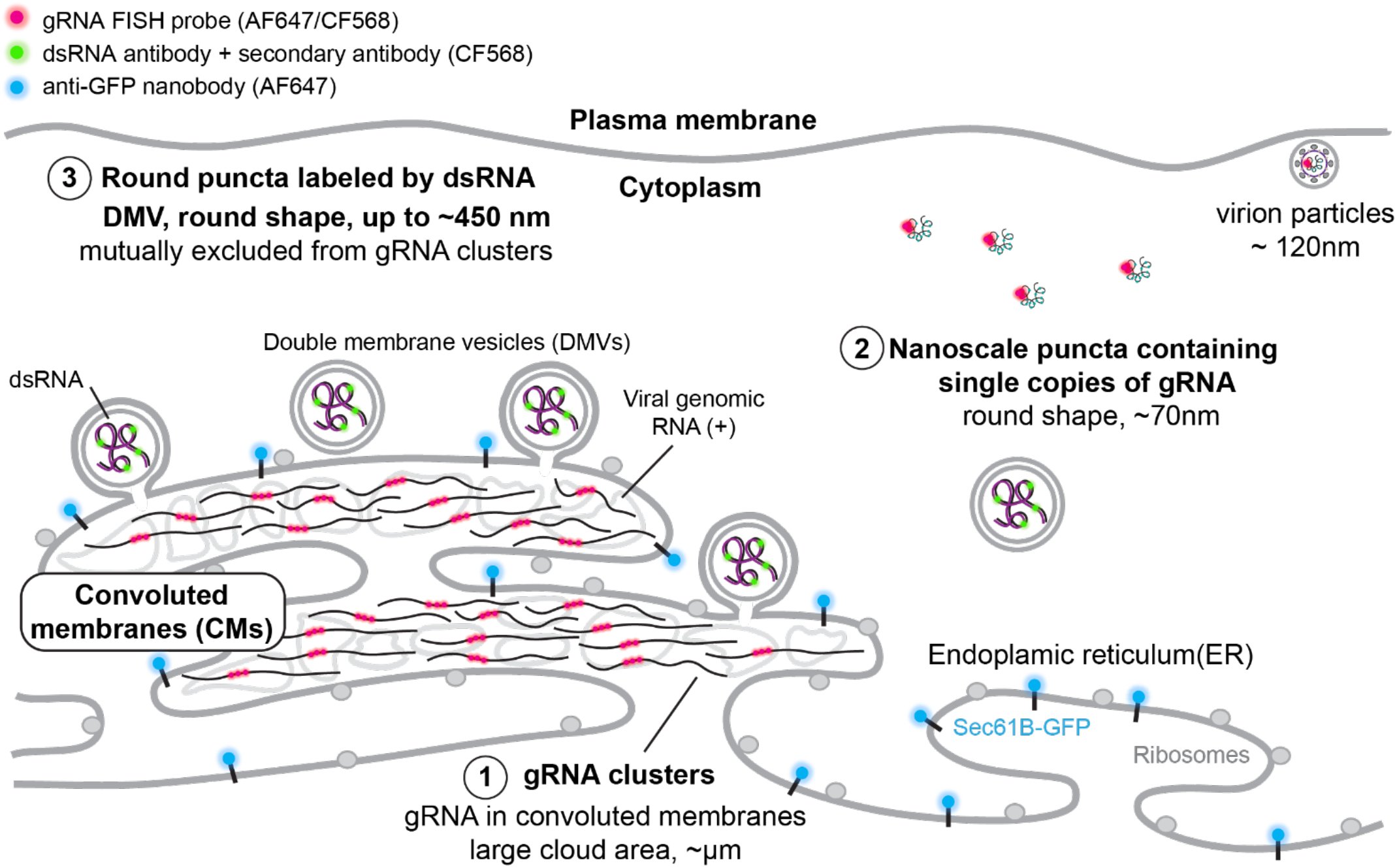
SR-based model of HCoV-229E gRNA and dsRNA distribution. A model showing the spatial organization of an HCoV-229E infected cell. The ER membrane is modified to create both convoluted membranes and DMVs. Large gRNA clusters were found in association with the ER membrane, whereas well-separated nanoscale gRNA puncta are often not connected to the ER. Rarely, assembled virions containing concentric colocalization of spike protein and gRNA are observed. The DMVs are filled with dsRNA to different extents. Some DMVs are still attached to the ER membrane, while others have budded off. These DMVs are separated from the convoluted membrane, where gRNA clusters appear to be the products of active replication.

First, our study shows that gRNA forms varying shaped irregular and extended web-like clusters that are often associated with the ER (**Figure 2C-F**). Several studies showed SARS-CoV and SARS-CoV-2 can modify the ER to create compartments such as double membrane vesicles and convoluted membranes (Goldsmith et al., 2004; Knoops *et al*., 2008; Snijder *et al*., 2006). The shape and the scale of the large gRNA clusters are reminiscent of the known convoluted membranes, which can contain up to hundreds of copies of the gRNA. As infection time increases, these clusters grow and can reach up to 9 square microns in area (**Figure 3L**). In contrast, at all stages of the infection, numerous well-separated nanoscale (~70 nm) gRNA puncta are present in the cell with a roughly constant size and brightness, which rarely associate with the ER membrane (**Figure 3A-C**). Since the brightness and size distribution of these nanoscale puncta match well with those for purified virions, these puncta appear to contain only a single copy of the gRNA. Importantly, 2-color SR imaging of purified virions shows that spike protein forms a concentric shell with the gRNA in the center (**Figure 3I**). As we rarely observe similar colocalization of gRNA puncta and spike protein (**Supplementary Figure 3D**) in the cell cytoplasm, the isolated gRNA puncta are likely to be free gRNA localized away from the ER. In addition, the sparsity of concentric colocalization suggests that intact virions are likely not present in the cell for very long. At the same time, separate gRNA and spike protein are scattered all over the remodeled ER membrane, but as is well-known, the spike protein is membrane-embedded possibly with other capsid proteins waiting for assembly, while the gRNA is cytoplasmic yet mostly confined inside the ER. It is necessary for the gRNA to transit to the other side of the ER enclosure to find the proper binding surface to assemble a virion. As gRNA as well as spike proteins are present in high abundance, the synthesis of these appears to be fast compared to virion assembly. The paucity of assembled virions suggests that assembled virions are likely exported readily and rapidly. Thus, we believe that the rate-limiting step is the cytoplasmic gRNA finding the nascent virion membrane studded with the required membrane-embedded proteins, a process which is dependent on diffusion as well as on the required RNA-binding proteins.

Occasionally, we observed nanoscale gRNA puncta at a higher density (**Figure 3A**, bottom left). As individual puncta are well separated, these could represent the vesicle packets (VP) observed in several EM studies which contain multiple virions. It is unclear whether the virions inside the packets are fully assembled virions or just single gRNA coated by nucleocapsid proteins (Knoops *et al*., 2008; Ogando *et al*., 2020).

Second, our SR reconstructions show that dsRNA (intermediates of RNA replication and transcription) forms larger, circularly shaped puncta of up to ~450 nm in diameter (**Figure 5E**). Several cryo-EM studies captured DMVs in CoV-infected cells from other organisms, the size of which ranged from 100 – 400 nm in diameter (Klein *et al*., 2020; Knoops *et al*., 2008; Wolff *et al*., 2020). We explored the hypothesis that our dsRNA puncta are located within DMVs. As infection time increases, the number of dsRNA puncta increases (**Figure 4D**), while the distribution of the size of dsRNA puncta stays the same. We note the circularity as well as the dimensions of these puncta (slightly smaller than expected DMV diameters). Thus, these dsRNA puncta are likely encapsulated by the lipid membrane of DMVs. Importantly, these dsRNA puncta do not colocalize with the ER (**Figure 2A-B**). As the maximum size of the dsRNA puncta is consistent with being within the lipid boundary of the DMVs, puncta that are over a couple hundred nanometers away from the ER signal likely are already severed from the rest of the ER (**Figure 5F-G and Figure 7**). At the same time, the dsRNA puncta adjacent to the ER could represent DMVs that are budding off from the ER membrane (**Figure 5F-G and Figure 7**). Interestingly, dsRNA puncta were not visualized by the ER membrane protein label we used in this study, suggesting that the composition and thus property of the viral manipulated membrane may be different from the rest of the ER, or the high radius of curvature of the DMVs or the membrane is too densely populated with viral proteins precluded the single-pass label native to the ER. This is based upon the commonly observed decrease in binding affinity as number of transmembrane passes decreases (Derganc and Copic, 2016; Larsen *et al*., 2020).

Third, we found that while nanoscale (~70 nm) gRNA puncta we identify as single copies of viral RNA typically have no dsRNA signal nearby, large gRNA clusters are decorated by dsRNA puncta at the periphery, without colocalization. This striking spatial separation in contrast to previous findings is likely caused by gRNA and dsRNA being stored in different compartments. One explanation for this observation is that the FISH probes cannot access the gRNA when it is hybridized in the dsRNA form either due to the base-pairing or competing binding of the dsRNA antibody. Furthermore, this might suggest the possibility that fully synthesized gRNA and the storage of its intermediate product (dsRNA) are separated, a potential distinction between the Alphacoronaviruses and Betacoronaviruses, in which DMVs are active sites of RNA synthesis (Knoops *et al*., 2008; Snijder *et al*., 2020). Another possibility is that the DMVs might only contain double-stranded sgRNAs (mRNA templates) that are not targeted by the FISH probes. The spatial separation of positive sense full-length gRNA and sgRNA synthesis might be programmed for optimal efficiency or in order to separate virion packaging and nonstructural protein expression. And lastly, it is also plausible that the dsRNA puncta represent the active replication sites, and the gRNA clusters represent replicated gRNAs that diffused away from the active replication sites.

Finally, we observed distinct responses of the gRNA clusters and dsRNA puncta with respect to Remdesivir treatment (**Figure 6**). Overall, Remdesivir reduces the confocal fluorescent signal from both gRNA and dsRNA (**Figure 6A**), as the ribonucleoside analog causes a reduction in viral RNA amplification (Beigel *et al*., 2020; Spinner *et al*., 2020; Wang *et al*., 2020). However, the drug treatment reduced the median size of gRNA clusters whereas the median size of dsRNA puncta moderately increased (**Figure 6F-G**). Combined with our observations that dsRNA and gRNA are spatially separated with radically different shapes, we propose that the active RNA replication happens within the convoluted membrane where gRNA forms clusters. Because the dsRNA objects (putative DMVs) did not change size upon drug treatment, DMVs, might be a temporary storage space for RNA replication intermediates before the next virion can be packaged. The benefit of such spatial organization is a subject for further investigation. It is unclear whether the storage of dsRNA has a particular benefit or is simply a byproduct, and the packaging of virions is slower than RNA replication. Clearly, there are many open questions concerning the release of RNA from DMVs for virion assembly or translation, and the pore structure found in other DMVs may be involved here (Wolff *et al*., 2020).

Our observations highlight the advantages of multicolor fluorescence SR imaging in studying the virus-host interaction with cellular context to observe nanoscale spatial organization of viral RNA and proteins in infected cells. The approach demonstrated here for the HCoV-229E-infected MRC5 cells can be applied to the investigation of other coronaviruses to elucidate the interactions of viral RNA with host cell organelles. In combination with high-throughput imaging systems, this method can aid in drug testing processes where assessment of RNA distribution is important. Broadly, we envision that future efforts will apply 3D advanced SR imaging techniques as well as the combination of single-molecule imaging and cryo-electron tomography to simultaneously achieve nanometer resolution and exquisite molecular specificity for virology research.

## ACKNOWLEDGEMENTS

This work was supported in part by the National Institute of General Medical Sciences Grant Nos. R35GM118067 (to W.E.M.) and the National Institutes of Health Common Fund 4D Nucleome Program No. U01 DK127405 (to L. S. Q.). We thank Lucy Shapiro for helpful advice, support, and careful review of the manuscript. We thank Maurice Youzong Lee (Genome Institute of Singapore) for helpful suggestions and assistance in clustering analysis. J.W. is a Mona M. Burgess Stanford Bio-X Fellow. We acknowledge use of the Biorender package for figure generation.

## METHODS

### Cell culture

The MRC-5 cells (Cat# CCL-171, ATCC) and HEK293T cells (Cat# CRL-3216, ATCC) were cultured in DMEM with GlutaMAX (Cat#10569010, Life Technologies) in 10% FBS (Cat#F0926, Sigma) at 37 °C and 5% CO_2_ in a humidified incubator.

### Lentivirus production

To produce lentivirus, HEK293T cells were cultured in 10-cm dishes and transiently transfected with 9 μg lentiviral plasmid pLV-ER-GFP (Cat# 80069, Addgene, a gift from Pantelis Tsoulfas), 8 μg pCMV-dR8.91, and 1 μg PMD2.G packaging plasmids using 25 μl TransIT^®^-LT1 Transfection Reagent (Cat# MIR 2306, Mirus). After 72 h of transfection, supernatant was filtered through 0.45 μm filters, concentrated using Lenti-X concentrator (Cat# VC100, ALSTEM) at 4°C overnight, and centrifuged at 1,500 × g for 30 min at 4°C to collect virus pellets. The virus pellets were resuspended in cold culture medium for storage at −80 °C for transduction of cells.

### Generation of stable cell line

To generate MRC-5-ER-GFP stable cells for ER imaging, 400 K MRC-5 cells were seeded in one well of a 6-well plate. One quarter of concentrated lentivirus expressing pLV-ER-GFP produced from one 10-cm dish of HEK293T cells were added to cells for incubation while seeding. After two days’ incubation, cells expressing GFP were sorted out using a SONY SH800S sorter. To clarify, these cells were only used for ER imaging; all other experiments used wild type cells.

### 229E production

Human coronavirus 229E (Cat# VR-740, ATCC) is amplified once by inoculating one T75 flask of fully confluency MRC-5 cells. The supernatant was collected at 48 h p.i. and centrifuged at 1500 g for 10 min to remove cell debris. The virus was aliquoted and stored at −80°C. The virus titer is determined by TCID50 assay.

### Infection of the cells by 229E

MRC-5 cells or MRC-5-ER-GFP cells were seeded at 4-4.5 × 10^4^ cells per well into poly-D-lysine-treated 8-well μ-slides (Cat# 80826, IBIDI) one day before infection. The cells were then infected with 229E at an MOI (Multiplicity of infection) of 0.1 or 0.2. At 6, 12, or 24 hpi, the cells were fixed for RNA FISH staining. For Remdesivir treatment, MRC5 cells were treated with 0.1 μM or 0.5 μM Remdesivir 30 minutes after 229E infection.

### Synthesis of the RNA FISH probes

Two sets of RNA FISH probes were designed targeting 229E, respectively, by utilizing the Stellaris^®^ RNA FISH Probe Designer (Biosearch Technologies, Inc., Petaluma, CA) available online at https://www.biosearchtech.com/stellaris-designer (version 4.2). The probe set targeting 229E (+) genomic RNA were designed targeting against the (+) RdRP-coding region within the ORF1a/b region of 229E, and the probe set targeting 229E total RNA (including genomic RNA and mRNA) were designed targeting the (+) N-protein-coding region and the 3’-UTR. A listing of all the FISH probes is provided at the end of the document. The designed probes were ordered with 5AmMC6 modifications from Integrated DNA Technologies in plate format of 25 nmole scale with standard desalting. Each probe was dissolved in water to a final concentration of 100 μM. The same set of probes was combined with equal volumes of each probe to get a stock of 100 μM mixed probes. The mixed probes were further desalted using ethanol precipitation. Briefly, 120 μL 100 μM probes were mixed with 12 μL 3 M sodium acetate (pH 5.5), followed by 400 μL ethanol. After precipitation at – 80°C overnight, probes were pelleted through centrifugation at 12,000 × g for 10 min at 4 °C, washed with precooled 70% (vol./vol.) ethanol for three times, air dried, and dissolved in water to make a 100 μM solution of probes. Then, 18 μL 100 μM probes was mixed with 2 μL 1 M NaHCO3 (pH 8.5), followed by 100 μg Alexa FluorTM 647 succinimidyl ester (NHS) (Cat# A37573, Invitrogen) or CF568, succinimidyl ester (NHS) (Cat# 92131, Biotium) dissolve in 2 μL dry DMSO (Cat# D12345, Invitrogen). The mixture was incubated for 3 days at 37 °C in the dark for conjugation and purified for 3 rounds using Monarch^®^ PCR & DNA Cleanup Kit (5 μg) (Cat# T1030S, NEB) following the manufacturer’s instructions. The estimated labeling efficiency of probes was calculated using the following equation:

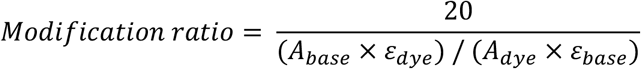

where ε_dye_ is 239,000 cm^-1^M^-1^, ε_base_ is 8,919 cm^-1^M^-1^, A_base_ is the absorbance of the nucleic acid at 260 nm, and Adye is the absorbance of the dye at 650 nm.

### RNA FISH

RNA FISH was performed following the Stellaris RNA FISH protocols available online (https://biosearchassets.blob.core.windows.net/assets/bit_stellaris_protocol_for_adherent_cells_in_96_well_glass_bottom_plates.pdf) (Femino et al., 1998; Raj et al., 2008). Briefly, cells cultured in 8-well μ-slides were washed with 1xPBS, fixed with 3.7% formaldehyde in 1xPBS at room temperature for 10 min, washed twice with 1xPBS, and then permeabilized with 70% (vol./vol.) ethanol for >1 hour at 4°C. After decanting 70% ethanol from wells, 200 μL Wash Buffer A (40 μL Stellaris^®^ RNA FISH Wash Buffer A (Cat# SMF-WA1-60, LGC Biosearch Technologies), 20 μL deionized formamide, 140 μL H2O) was added to cells and incubated at room temperature for 5 min. After decanting Wash Buffer A, 100 μL Hybridization Buffer (90 μL Stellaris^®^ RNA FISH Hybridization Buffer (Cat# SMF-HB1-10, LGC Biosearch Technologies), 10 μL deionized formamide) containing 2 μL 12.5 μM RNA FISH probes was added into each well and incubated for 16 hours at 37 °C in the dark. Then cells were washed with Wash Buffer A for 30 min at 37°C in the dark, washed with Wash Buffer A containing DAPI for 30 min at 37°C in the dark, and stored in Wash Buffer B (Cat# SMF-WA1-60, LGC Biosearch Technologies) for imaging. DAPI was only added to the samples for DL imaging and not added to the samples for SR imaging.

For two-color RNA FISH staining, 2 μL 12.5 μM AF647-labeled RNA FISH probes targeting the RdRp region and 2 μL 12.5 μM CF568-labeled RNA FISH probes targeting the N-protein and 3’-UTR region were added together into 100 μL Hybridization Buffer for incubation in each well. The other steps of RNA FISH staining are the same.

### Simultaneous RNA FISH and immunofluorescence staining

Cells cultured in 8-well μ-slides were fixed and permeabilized as described for RNA FISH. All following steps were performed at 37 °C. After decanting 70% ethanol, cells were washed once with Wash Buffer A at room temperature for 5 min. For simultaneous staining of gRNA and dsRNA, wild type cells were incubated with 100 μL Hybridization Buffer containing 2 μL 12.5 μM AF647-labled gRNA FISH probes and 1:1000 anti-dsRNA antibodies (Scicons J2 mouse monoclonal antibody) for 4 hours in the dark. Then cells were incubated with 1:20000 CF™ 568 Donkey anti-mouse antibody (Sigma, SAB4600075) for 30 min, washed with Wash Buffer A containing DAPI for 30 min in the dark, and stored in Wash Buffer B for imaging. For simultaneous staining of gRNA and ER membrane, transduced cells were incubated with 100 μL Hybridization Buffer containing 2 μL 12.5 μM CF568-labled gRNA FISH probes and 1:2000 AF647-labeled anti-GFP nanobody (Chromotek, gb2AF647-50) for 4 hours in the dark. Then cells were washed twice with Wash buffer A containing DAPI for 30 min in the dark and stored in Wash Buffer B for imaging. For simultaneous staining of dsRNA and ER membrane, transduced cells were incubated with 100 μL Hybridization Buffer containing 1:1000 anti-dsRNA antibodies and 1:2000 AF647-labeled anti-GFP nanobody for 4 hours in the dark. Then cells were incubated with 1:20000 CF™ 568 Donkey anti-mouse antibody for 30 min, washed with Wash Buffer A containing DAPI for 30 min in the dark, and stored in Wash Buffer B for imaging. For simultaneous staining of gRNA and Spike proteins, wild type cells were incubated with 100 μL Hybridization Buffer containing 2 μL 12.5 μM CF568-labled gRNA FISH probes and 1:500 anti-Spike antibodies (Cat# PAB21477-500, The Native Antigen Company) for 4 hours in the dark. Then cells were incubated with 1:1000 AF647-labeled Donkey anti-Rabbit antibody (Cat# A-31573, Invitrogen) for 30 min, washed with Wash Buffer A containing DAPI for 30 min in the dark, and stored in Wash Buffer B for imaging. DAPI was not added to the samples for SR imaging.

### RNA FISH and immunofluorescence staining of purified virions

4×10^4^ TCID50 of 229E virus in 200 μl DMEM medium were added into one well of poly-D-lysine-treated 8-well μ-slides and incubated at 4°C for 24 h to coat the virions onto the surface of the well. Then, the media containing virions was removed and 3.7% formaldehyde in 1xPBS was directly added to the well for a 10-min incubation at room temperature. The well coated with virions was washed twice with 1xPBS, permeabilized with 70% (vol./vol.) ethanol for >1 hour at 4°C, and washed with Wash Buffer A at room temperature for 5 min. After decanting Wash Buffer A, 100 μL Hybridization Buffer containing 2 μL 12.5 μM RNA FISH probes was added to the well and incubated for 16 hours at 37 °C in the dark. The wells coated with virions were washed twice with Wash Buffer A by incubating for 30 min at 37 °C in the dark and stored in Wash Buffer B for imaging.

For two-color staining of purified 229E virions with both gRNA FISH probes and anti-spike antibody, the virions coated on poly-D-lysine-treated 8-well μ-slides were fixed, permeabilized, and washed with Wash Buffer A as described in the previous paragraph. After decanting Wash Buffer A, 100 μL Hybridization Buffer containing 2 μL 12.5 μM CF568-labeled gRNA FISH probes was added into the well coated with virions and incubated for 14 hours at 37 °C in the dark. Then, 1:500 antispike antibodies were added to the hybridization system containing FISH probes for another 4-h incubation at 37 °C in the dark. Virions were next incubated with 1:1000 AF647-labeled Donkey antiRabbit antibody in Wash Buffer A for 30 min at 37 °C in the dark, washed with Wash Buffer A for 30 min at 37 °C in the dark, and stored in Wash Buffer B for imaging.

### Spinning disk confocal microscopy

Confocal microscopy was performed on a Nikon TiE inverted spinning disk confocal microscope (SDCM) equipped with a Photometrics Prime 95B camera, a CSU-X1 confocal scanner unit with microlenses, and 405 nm, 488 nm, 561 nm, and 642 nm lasers, using the 60 × PLAN APO IR water objective (NA = 1.27). Images were taken using NIS Elements version 4.60 software with Z stacks at 0.3 μm steps. The camera pixel size of SDCM is 0.183 μm/pixel. The pinhole size is 50 μm. Only one Z slice is used for all images shown.

### Confocal image analysis

To quantify the number of infected MRC5 cells at different stages of the infection in **Fig 4C** and **Fig 6A**, we counted cells with both gRNA and dsRNA staining as infected cells in FIJI (Schindelin et al., 2012). The infected cells were further characterized manually into two types in **Fig 4D**: type I shows scattered dot-like localization of both gRNA and dsRNA; type II shows large clusters of gRNA decorated with small dot-like dsRNA puncta. The plots showing the number of different cell types and total cell number were generated by Prism 9.

To quantify the number of dsRNA puncta in each cell in **Fig 4D** and **Fig 6B**, the Trackmate (Tinevez et al., 2017) plugin in FIJI was used. We experimentally tested parameters to detect dsRNA puncta using LoG detector in the Trackmate plugin, and the same parameters were used for all conditions (estimate blob diameter=0.5; Threshold=50, with median filter and sub-pixel localization). The Trackmate analysis was performed on full 3D stacks. The plots showing the number of detected puncta were generated by Prism 9.

### Super-resolution microscopy

To accurately determine the location of the various two-color labeled targets in the cell, we use super-localization to pinpoint the location of single molecules where photobleaching was used to reduce the emitting concentration. Cells were cultured and fixed as previously described. Before SR imaging, the PBS was replaced by a reducing and oxygen-scavenging buffer (Halpern *et al*., 2015) optimized for dSTORM blinking that consists of 100 mM Tris-HCl (Invitrogen), 10% (wt/vol) glucose (BD Difco), 2 μl/ml catalase (C100, Sigma-Aldrich), 560 μg/ml glucose oxidase (Sigma-Aldrich), and 50 mM cysteamine (Sigma-Aldrich). For two-color imaging, the buffer was also supplemented with 71.5 mM β-mercaptoethanol (Sigma-Aldrich).

Single-molecule imaging experiments were performed on a custom epifluorescence microscope (Nikon Diaphot 200) equipped with a Si EMCCD camera (Andor iXon DU-897) and a high NA oil-immersion objective (Olympus UPlanSapo 100x/1.4 NA). Molecules labeled with AF647 were excited with 642-nm, 1-W continuous-wave laser (MPB Communications Inc.) at ~ 6.5 kW/cm^2^ whereas Molecules labeled with CF568 were excited with 561 nm 200 mW continuous-wave laser 10 kW/cm^2^. An exposure time of 50 ms and a calibrated EM gain of 193 was used for image acquisition. The emission from fluorescent molecules was collected through a 4-pass dichroic mirror (Semrock, Di01-R405/488/561/635) and filtered by a ZET561NF notch filter (Chroma), ZET642NF notch filter (Chroma), a 561 EdgeBasic long-pass filter (Semrock), and ET610/60 bandpass filter (Chroma) for CF568 detection or ET700/75m bandpass filter (Chroma) for AF647 detection. For two-color experiments, AF647 data was acquired before CF568 data to avoid bleaching of AF647 via 561 nm excitation.

### SR data analysis

We processed approximately 40000 frames of single-molecule images for 2D Gaussian fitting by ThunderSTORM (FIJI) (Schindelin *et al*., 2012; Tinevez *et al*., 2017). We used the local maximum algorithm to estimate localization of molecules with a 3xstd peak intensity threshold and 8-neighborhood connectivity. A 3-pixel fitting radius was used to fit the point spread function using weighted least square method. Localization precision of AF647 and CF568 was determined experimentally to be around 10 nm. Dye conjugated FISH probes were immobilized to the cell chamber and imaged under the same condition as cellular imaging. The localization precision is estimated by taking the standard deviation of the multiple measured positions of the immobile single molecules (**Supplementary Figure 2F**). Sample drift was corrected by cross-correlation, followed by filtering (sigma < 200 nm; uncertainty of localization < 20 nm). To suppress biases from overcounting, blinking events were merged if occurred within 30 frames and 20 nm. Images were reconstructed as 2D histograms with a bin size of 23.4 nm, corresponding to a five-time magnification (camera pixel size 117.2 nm).

For two-color SR imaging, the two channels were registered by imaging 200 nm TetraSpeck beads (Invitrogen) visible in both channels, followed by affine transformation using MATLAB’s built-in function fitgeotrans (fiducial registration error ~ 7 nm). The calculated transformation matrix was then applied to the respective reconstructed images.

### Virion quantification

For every field of view of purified virions, a diffraction-limited image was acquired before single-molecule data acquisition. In the DL image, each virion labeled with gRNA is fitted with a 2D gaussian function (the same way as described in SR data analysis) and the total intensity of each virion can be extracted. Next, the integrated brightness of single molecules (each FISH probe) within the virions is measured by fitting a 2D Gaussian function. An average value of the brightness of each individual FISH probe was determined. Finally, the number of FISH probes labeling each virion is calculated by dividing the total brightness of each virion by the brightness of individual FISH probes. For spike labeled virions, the shapes were often more elliptical than circular. Thus, the virions here were fit with an elliptical 2D Gaussian function. The average of the major and minor axes was used to calculate an estimation for the full width at half maximum of the virion.

### Clustering analysis

To calculate the sizes of gRNA and dsRNA clusters as well as the number of molecules within each cluster, we used Voronoi tessellation on the single molecules from the previous step using a custom MATLAB script (**Supplementary code.A**). The Voronoi cell was determined for each individual molecule by MATLAB’s built-in function “Voronoi”. For a molecule to be considered within a cluster, we used 500 nm^2^ and 1000 nm^2^ as cell area threshold for gRNA and dsRNA molecules, respectively. We checked for robustness against the choice of these thresholds and could verify that changing their values does not dramatically affect cluster sizes or numbers. All Voronoi cells that share borders were then merged into a single cluster (**Supplementary Figure 4 H-K**). After merging Voronoi cells, peripheral molecules that had much larger cell areas were added to a given cluster if their distance to the nearest neighbor located within that cluster was smaller than the shortest pairwise distance within the cluster. Finally, the area of the clusters was calculated by the summation of all individual cell areas of the cluster (excluding the peripheral molecules).

